# Characterization of Alternative Splicing During Mammalian Brain Development Reveals the Magnitude of Isoform Diversity and its Effects on Protein Conformational Changes

**DOI:** 10.1101/2023.10.11.561865

**Authors:** Leila Haj Abdullah Alieh, Beatriz Cardoso de Toledo, Anna Hadarovich, Agnes Toth-Petroczy, Federico Calegari

**Affiliations:** CRTD-Center for Regenerative Therapies Dresden, School of Medicine, TU Dresden, Germany; Max Planck Institute of Molecular Cell Biology and Genetics, 01307 Dresden, Germany; Center for Systems Biology Dresden, 01307 Dresden, Germany; Cluster of Excellence Physics of Life, TU Dresden, 01062 Dresden, Germany; Brain Mind Institute, Faculty of Life Sciences, École Polytechnique Fédérale de Lausanne, Switzerland

**Keywords:** Alternative splicing, brain development, neurogenesis, protein structure prediction, isoform- specific protein conformations

## Abstract

Regulation of gene expression is critical for fate commitment of stem and progenitor cells during tissue formation. In the context of mammalian brain development, a plethora of studies have described how changes in the expression of individual genes characterize cell types across ontogeny and phylogeny. However, little attention was paid to the fact that different transcripts can arise from any given gene through alternative splicing (AS). Considered a key mechanism expanding transcriptome diversity during evolution, assessing the full potential of AS on isoform diversity and protein function has been notoriously difficult. Here we capitalize on the use of a validated reporter mouse line to isolate neural stem cells, neurogenic progenitors and neurons during corticogenesis and combine the use of short- and long-read sequencing to reconstruct the full transcriptome diversity characterizing neurogenic commitment. Extending available transcriptional profiles of the mammalian brain by nearly 50,000 new isoforms, we found that neurogenic commitment is characterized by a progressive increase in exon inclusion resulting in the profound remodeling of the transcriptional profile of specific cortical cell types. Most importantly, we computationally infer the biological significance of AS on protein structure by using AlphaFold2 and revealing how radical protein conformational changes can arise from subtle changes in isoforms sequence. Together, our study reveals that AS has a greater potential to impact protein diversity and function than previously thought independently from changes in gene expression.

## INTRODUCTION

The lack of correlation between genes number and complexity of eukaryotic organisms has puzzled biologists for decades (Hahn and Wray, 2002; Thomas, 1971). As one example, the worm *C. elegans* shares with us humans a comparable number of protein-coding genes despite major differences in tissue diversity and organ complexity (Cunningham *et al*., 2022). The reasons for such a lack of correlation are not entirely clear but an evolutionary increase in abundance and complexity of alternative splicing (AS) can partly explain this discrepancy. In fact, changes in AS, transcriptional initiation and 3′ cleavage/polyadenylation sites have great potential to increase transcripts diversity from a common pool of genes and, thus, considerably expand the genomic coding potential (Nilsen and Graveley, 2010; Reixachs-Solé and Eyras, 2022). Consistently, frequency and heterogeneity of AS have expanded across evolution (Schaefke *et al*., 2018; Verta and Jacobs, 2022) and positively correlate with organism complexity (Chen *et al*., 2014; Yang *et al*., 2021). Although its full extent and biological significance remain poorly investigated (Pickrell *et al*., 2010; Saudemont *et al*., 2017; Tress *et al*., 2017), AS is thought to have enormous potential to redefine proteins sequence and, as a result, modulate their function by changing their stability, localization, interaction with other molecules and so forth (Kelemen *et al*., 2013; Marasco and Kornblihtt, 2023).

AS profiles that are highly specific not only across species but also across organs within the same species result from the concerted modulation of AS events (i.e.: inclusion/exclusion of exons, parts of exons or intron retention) (Barbosa-Morais *et al*., 2012; Buljan *et al*., 2012; Ellis *et al*., 2012; Rodriguez and Pozo, 2020). Notably, of different organs, the brain is among the ones with the highest proportion of spliced genes (Mazin *et al*., 2021; Yeo *et al*., 2004) and highly specific and conserved AS profiles, including microexons (≤ 27 nt) (Irimia *et al*., 2014; Raj and Blencowe, 2015). One potential reason for the prevalence of AS in the brain is the extreme diversity of cell types characterizing this organ and arising during development from neural stem cells (NSC). More specifically, during embryonic development NSC initially undergo proliferative division to expand their pool and later switch to differentiative division to generate neurogenic progenitors (NP) giving rise to neurons (N) specifying into hundreds of different neuronal cell types critical for brain function (Villalba *et al*., 2021). While several studies have investigated the role of AS in the context of neuronal maturation and specification (Furlanis and Scheiffele, 2018; Raj and Blencowe, 2015; Weyn-Vanhentenryck *et al*., 2018; Zhang *et al*., 2016) and linked neural-specific splicing factors to neurodevelopmental disorders (Chau *et al*., 2021; Quesnel-Vallières *et al*., 2016), relatively little attention has focused on the role of AS in neurogenic commitment and, specifically, cell fate transition from NSC to NP. Limited examples among these studies include the identification of neurogenic-specific splicing factors (Han *et al*., 2022; Verdile *et al*., 2023), splice site mutations arising in evolution and resulting in progenitor expansion (Florio *et al*., 2016), and crosstalk between epigenetic and splicing programs (Sahu *et al*., 2022). Despite of this, a comprehensive annotation of isoform diversity across neuronal cell types and assessment of its potential effects on protein structure are missing.

Several technical limitations contribute to make the study of AS in the context of cell fate commitment particularly challenging. For example, assessment of AS traditionally relies on annotation databases that are often neither organ- nor cell-specific and usually neglect underrepresented cell sub-populations (Morillon and Gautheret, 2019; Zhang *et al*., 2020). While the advent of long read sequencing (LRS) has significantly improved the identification of full-length and new transcripts including novel exons and cryptic splice sites(An *et al*., 2018; Oikonomopoulos *et al*., 2020), a reliable assessment of isoform abundance and relative proportion of splice variants remains unfeasible with this method (Sarantopoulou *et al*., 2021). Conversely, quantification of relative exon inclusion relies on the use of short read sequencing (SRS) that, on the other hand, does not allow the reconstruction of full-isoform variants. Independently from the use of LRS or SRS, additional technical aspects are to be considered in the use of single-cell sequencing data to assess exon inclusion such as, among others, biases introduced by PCR amplification, degree of RNA coverage, dropouts rates and complexity of computational analysis (Arzalluz-Luque and Conesa, 2018; Buen Abad Najar *et al*., 2020; Westoby *et al*., 2020). While great effort is being made to overcome these limitations, capturing cell type-specific AS dynamics that is both quantitative and comprehensive of full-length transcript information currently requires combination of both SRS and LRS performed in parallel on the same cell pool. This was seldom attempted (Gupta *et al*., 2018; Joglekar *et al*., 2021) and, to the best of our knowledge, never for specific cell types of the developing mammalian brain. Even more limiting, systematic assessment of the consequences of AS on protein structure and putative function in cell fate commitment is entirely lacking.

To assess AS dynamics and explore its significance during brain development, we took advantage of an extensively characterized double-reporter mouse model that allows the identification of NSC, NP and N during corticogenesis by their combinatorial expression of RFP and/or GFP (Aprea *et al*., 2013, 2015). This dual-reporter mouse line was instrumental for the characterization of the transcriptome and epigenome defining neurogenic commitment including the functional assessment of pioneer transcription factors, different classes of non-coding RNAs (long/circular/micro) and DNA methylation (Aprea *et al*., 2013, 2015; Artegiani *et al*., 2015; Dori *et al*., 2019, 2020; Mestres and Calegari, 2023; Noack *et al*., 2019). Combining the use of this dual-reporter mouse line with LRS and SRS and bioinformatic tools, we here describe a novel workflow allowing us to reconstruct cell type-specific transcriptome diversity during brain development and quantitatively assess AS events. By this, we describe nearly 50,000 new transcripts including novel exons, splice sites and/or microexons, thus, uncovering the full spectrum of splicing dynamics accompanying fate transitions from NSC to NP and N.

Next, by using AlphaFold2 (Jumper *et al*., 2021; Varadi *et al*., 2022) we additionally inferred the biological significance of several observed AS events on the resulting protein sequence and structure by modeling the 3D conformational changes resulting from isoform switching characterizing specific cell states. Remarkably, this highlighted that nearly 40% of isoform pairs originating from the same gene exhibited large global conformational changes including fold switches. In addition, we describe the occurrence of regions with identical sequence yet adopting profoundly different secondary structures (secondary structure element switches), such as alpha-helix vs. beta-sheet, depending on distant AS events, thus, revealing that even negligible changes in exon usage can induce large conformational changes influencing the functional properties of proteins.

Overall, our study combines the use of a validated mouse model with new sequencing annotation, computational pipelines and advanced machine learning-based protein modeling in order to provide a comprehensive assessment of cell-type specific AS profiles and its potential biological significance in mammalian brain development.

## RESULTS

### A new cell type-specific transcriptome assembly of the developmental mouse cortex

We began our study by reconstructing a new cell type-specific transcriptome of the developing mouse cortex. To this aim, we assembled previously generated SRS data of NSC, NP and N at the peak of neurogenesis at mouse embryonic day (E) 14.5 (Aprea *et al*., 2013, 2015). Using Hisat2-Stringtie (Kim *et al*., 2019; Pertea *et al*., 2015), this resulted in 25,742 transcripts from 11,952 genes (Fig. 1A; left). Such ratio of about 2:1 transcripts-to-genes was relatively low considering the 5:1 ratio previously reported in mouse (Cunningham *et al*., 2022) and likely resulting from intrinsic limitations in the reconstruction of full-length transcripts from SRS rarely spanning across multiple exons and often failing to resolve ambiguities at complex loci (Shumate *et al*., 2022).

**Figure 1.**
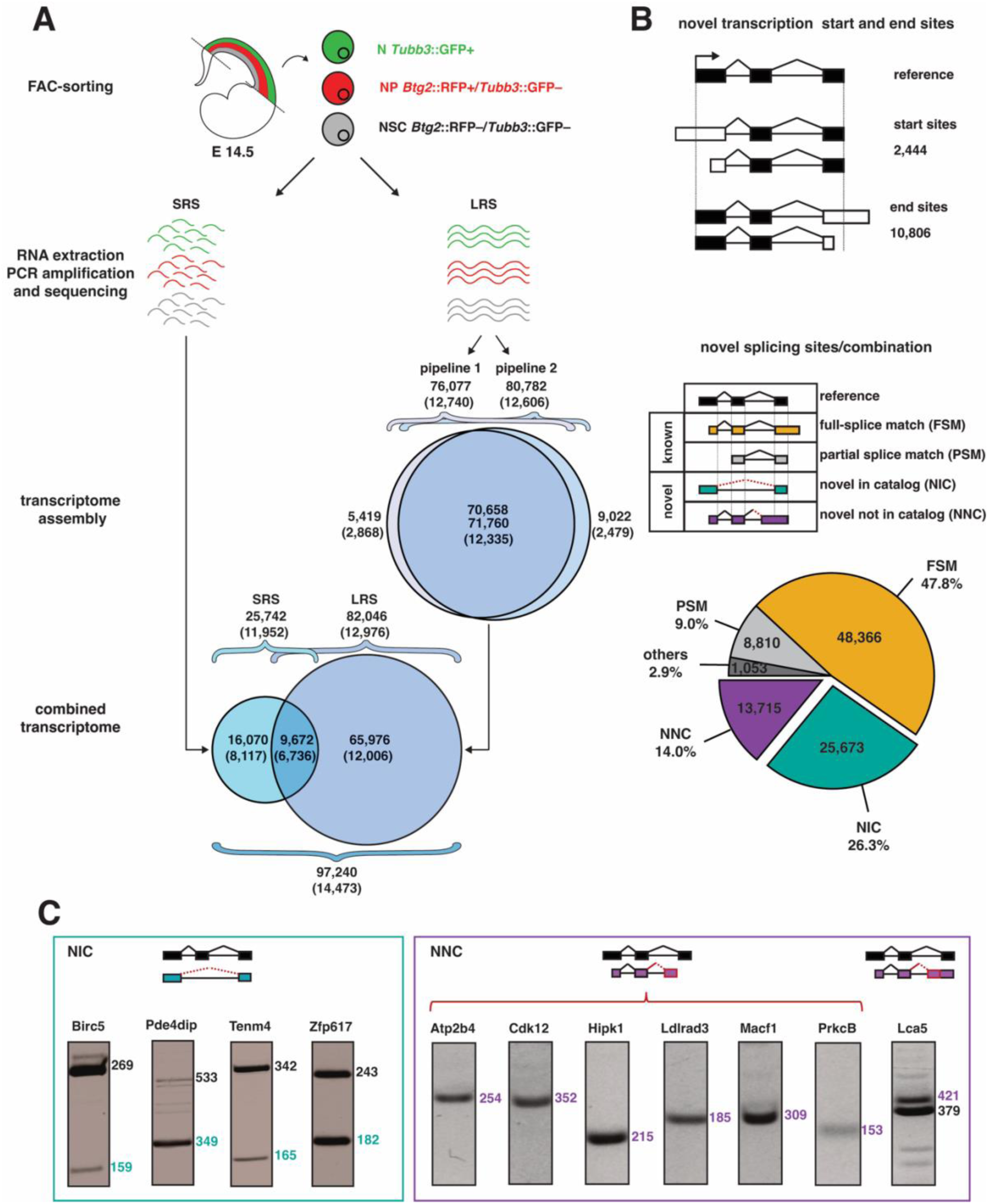
Transcriptome assembly of NSC, NP and N. **A)** Drawing exemplifying the combination of SRS and LRS datasets from cortical cell types obtained from E14.5 Btg2::RFP/Tubb3::GFP mouse embryos as previously described (Aprea et al., 2013; 2015). Upon SRS and LRS (left and right, respectively) assembling of transcripts (genes within parentheses) was performed using Hisat2-Stringtie (SRS) or two alternative pipelines (LRS) that were merged into a combined assembly of 97,240 transcripts from 14,473 genes. **B)** Classification of known and novel transcripts upon splice junction assembly obtained comparing either transcripts start/end sites (top) or internal splice junction (bottom). The latter included known transcripts whose splice junctions (continuous lines) either completely or partially matched reference junctions (full- or partial-splice match, FSM or PMS, yellow or gray, respectively), and novel transcripts whose splice junctions (dotted lines) either matched novel combinations of known junctions (novel in catalog, NIC, green) or were never described before (novel not in catalog, NNC, purple). **C)** Validation of novel splice junctions (red) resulting in one RT-PCR product when involving a novel first or last exon and in two bands when involving a novel internal exon (i.e. in Lca5).

Given these limitations of SRS, we next exploited the use of LRS to expand our assessment of transcriptome diversity. To this aim, we extracted and sequenced mRNA from NSC, NP and N from the E14.5 mouse cortex as previously described (Fig. 1A; top) (Aprea *et al*., 2013, 2015). PacBio high-quality transcripts (accuracy ≥ 99%) were next processed by two alternative bioinformatic pipelines (pipeline 1 and 2, see Material and Methods) for quality control and removal of redundant or false-positive transcripts and artifacts. This resulted in 76,077 and 80,782 transcripts (12,740 and 12,606 genes) with about 90% overlap in both pipelines and 5,419 and 9,022 transcripts being only detected by pipeline 1 or 2, respectively (Fig. 1A; right). Importantly, the vast majority of 5’ (> 85%) and 3’ (> 95%) ends of these pipeline-specific transcripts were supported by Ensembl coordinates, data from CAGE and poly(A) motifs and/or previous reports (Veiga *et al*., 2022) and therefore unlikely to be the product of degradation. For these reasons, we decided to merge all LRS data into a unique assembly of 82,046 transcripts (12,976 genes) and further combine these with the 16,070 transcripts predicted from SRS data that were not already included in the LRS dataset resulting in a novel transcriptome assembly of 97,240 transcripts (14,473 genes) (Fig. 1A; bottom). With a new transcripts-to-genes approaching the 7:1 ratio, this exceeded by almost 2-fold the magnitude of transcript diversity recently described in the adult brain (Leung *et al*., 2021). This is remarkable when considering that the E14.5 mouse brain is expected to contain a much lesser degree of cell diversity than the adult brain.

In addition to 2,444 and 10,806 transcripts with novel transcription start or end site, respectively, we found that 40% of transcripts in our assembly were not annotated in any database and were therefore novel either due to the presence of new splice sites (novel not in catalog) or splice sites that are known but combined in new ways (novel in catalog) (13,715 and 25,673 transcripts from 5,392 and 6,635 genes, i.e.: 14% and 26%, respectively) (Fig. 1B). We next sought to validate the existence of novel transcripts belonging to both categories. To this aim, we selected candidate novel splice junctions by several criteria including: i) coverage by ≥ 30 junction reads, ii) low variability among replicates (coefficient of variation < 0.5), iii) unique new junction per gene, and iv) belonging to the top 85% most expressed genes. This resulted in a subset of 471 candidates of which only about 140 (30%) were predicted to give a maximum of 2 PCR products and, thus, be suitable candidates for validation by RT-PCR. Among these 140, we finally selected 11 for validation, which was successful in all cases (Fig. 1C).

Taken together, our analysis provides the most complete cell type-specific assembly of full-length transcriptomes of the developing mammalian cortex to date including nearly 50,000 potentially new transcripts.

### Patterns of AS and inclusion events characterizing neurogenic commitment

Next, we assessed AS characterizing the transition from NSC to NP and N. Quantitative assessment of isoforms abundance is notoriously difficult and unreliable both in the analysis of SRS as well as LRS data (Sarantopoulou *et al*., 2021). Therefore, we focused our study on classifying splicing events, namely cassette exons, alternative donor or acceptor splice sites and intron retention (Marasco and Kornblihtt, 2023) (Fig. 2A). For this, we used our full transcriptome assembly as a reference while quantifying splicing events based on SRS data only by using Whippet (Sterne-Weiler *et al*., 2018). Level of inclusion of a splicing event was then expressed as percentage spliced in (PSI), and considered differentially spliced based on the commonly used threshold of ≥ 10% with a high prediction confidence (see Material and Methods).

**Figure 2.**
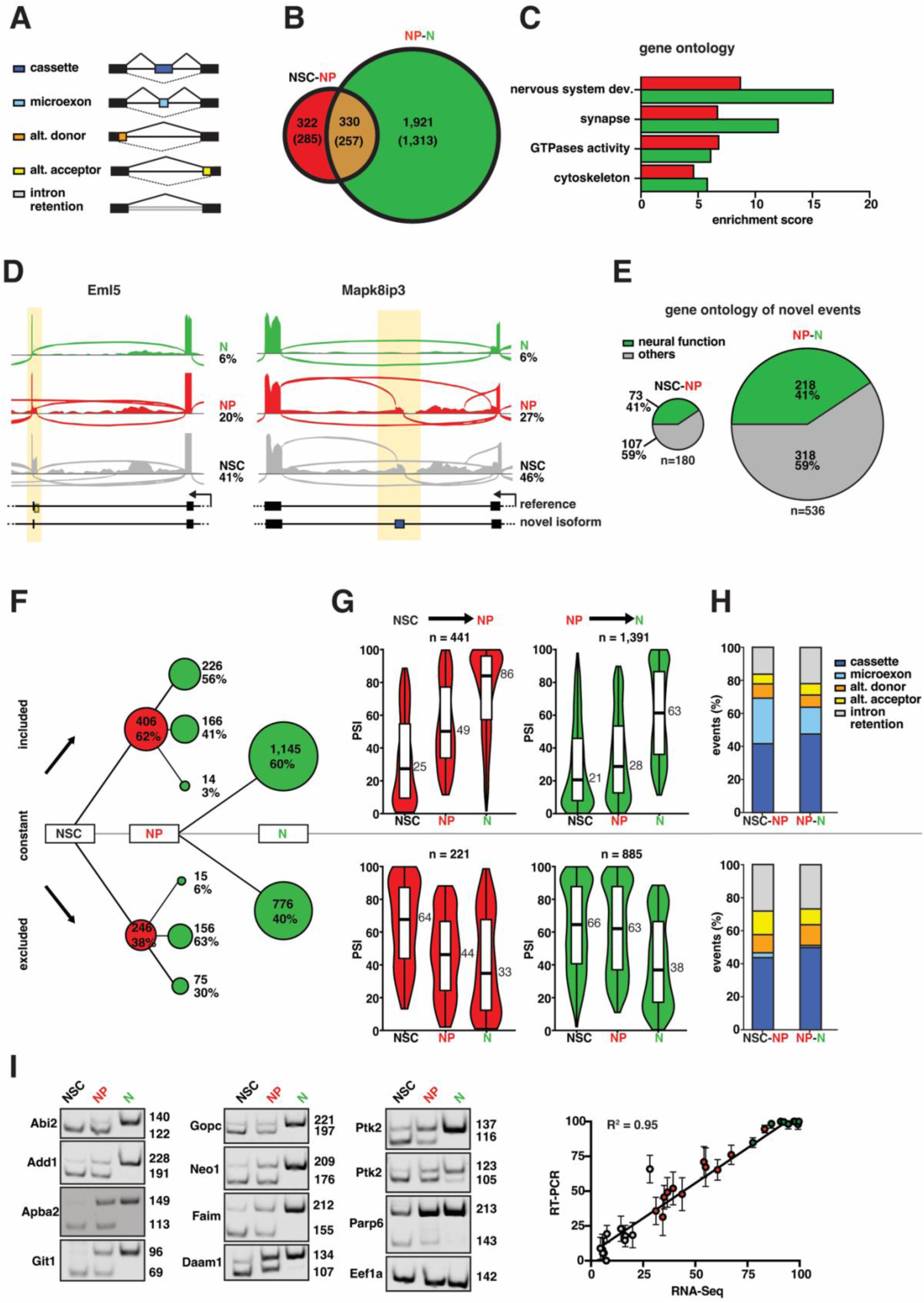
Assessment of AS during neurogenic commitment. **A**) Classification of types of splicing events into: cassette exons, microexons, alternative donor, alternative acceptor and intron retention. **B-C**) Diagrams showing the number of differentially spliced events and corresponding genes (parentheses) (**B**) and enrichment scores for their gene ontology annotation (**C**). Analyses were performed for differentially spliced events specifically in the NSC-NP (red), NP-N (green) transitions (**B** and **C**), or both (**B**, overlapping area). **D**) Sashimi plots of AS events involving novel transcripts. **E**) Proportion of novel transcripts detected in our study to undergo AS and known to regulate nervous system development or function. **F**) Patterns of AS classified as events being included, excluded, or unchanged (lines pointing up, down, or flat, respectively) in the progression from NSC to NP and NP to N (red and green circles, respectively). Area of circles are in scale to the number of events. Note the green circles stemming out of the red circles and representing the pattern of AS events changing in consecutive cellular transitions. **G**) Violin plots representing the PSI assessed in NSC, NP and N of events differentially included (top) or excluded (bottom) in the transition from NSC to NP (red, left) or NP to N (green, right). Median PSI values are indicated at the right of each violin plot. Note the gradual and constant changes (left), as opposed to the N-specific, single step change (right), occurring in the NSC-NP vs. NP-N transitions. **H**) Relative proportion of types (see legend and figure A) of included (top) or excluded (bottom) splicing events. Note the abundance of microexon inclusion (light blue; top) virtually undetected among excluded events (bottom). **I**) Validation of inclusion events using primers flanking the alternative exon and giving one exclusion (lower) and one inclusion (higher) qRT-PCR band (left) whose quantification showed a very high correlation coefficient (R^2^ = 0.95, p < 0.0001) (right) when compared to our bioinformatic analysis of combined SRS and LRS transcriptomes of cell types (colors).

We found a total of 2,573 splicing events (1,643 genes) changing their inclusion in subsequent stages of the neurogenic lineage, with 652 events (518 genes) differentially spliced during the transition from NSC to NP and 2,251 events (1,463 genes) from NP to N among which 330 events (257 genes) changed inclusion in both NSC-NP and NP-N transitions (Fig. 2B). Consistently with the assessment of neurogenic commitment, gene ontology analysis of all differentially spliced genes highlighted an enrichment in terms related to neurogenesis and *nervous system development* in both NSC-NP and NP-N (enrichment scores: 8.7 and 16.8, respectively) with similarly high enrichment scores related to *synapse*, *GTPase* and *cytoskeleton* also found (Fig. 2C). Several features were detected among these differentially spliced isoforms such as inclusion of novel sequences or use of novel splice junctions (examples in Fig. 2D) with a significant proportion of them (41%) being associated with gene ontology terms related to *neural function* (73/180 genes in NSC-NP and 218/536 genes in NP-N) (Fig. 2E). In turn, these findings highlight the extent of AS within the neurogenic lineage underscoring its potential to regulate corticogenesis to a much greater degree than previously appreciated.

Next, to reveal the dynamics of AS during neurogenic commitment we assessed PSI of variants characterizing NSC, NP and N finding a considerable overrepresentation of inclusion (62%) relative to exclusion (38%) events. This held true in both transitions from NSC to NP (406 vs 246 events) and NP to N (1,386 vs 865 events) (Fig. 2F). Moreover, we observed that the overwhelming majority of inclusion/exclusion events in the transition from NSC to NP continued their trend of inclusion/exclusion, or remained constant, in the following transition from NP to N while only a negligible fraction of events showed contrasting patterns, namely included in NP but excluded in N (14 events, 3%) or vice versa (15 events, 6%) (Fig. 2 F). In turn, such predominance of inclusion events, and bias in splicing patterns, during development are in agreement with previous reports not only in the mouse brain (Irimia *et al*., 2014; Weyn-Vanhentenryck *et al*., 2018) but also in other organs and species, from chicken to opossum and primates, including humans (Mazin *et al*., 2021).

Categorizing splicing events according to their patterns of inclusion/exclusion provided a first glance over the trends of AS. To achieve a more comprehensive view, we next expanded our previous assessment of the number and relative abundance of inclusion/exclusion events (Fig. 2F) to also consider their magnitude, i.e. their change in PSI (Fig. 2G). We first started by considering differentially spliced exons in the transition from NSC to NP. Among this group, the PSI of exons gaining inclusion (441) constantly increased from a median of 25 in NSC to 49 in NP and 86 in N (Fig. 2G; red, top-left). These values indicate that, overall, events that started to gain inclusion during neurogenic commitment occurred within the least abundant isoforms of NSC and that these isoforms became the most abundant in N. Conversely, exons undergoing exclusion from NSC to NP (221) belonged to the predominant isoforms in NSC that constantly decreased their median PSI from 64 to 44 in NP and 33 in N, ultimately becoming the least-represented isoforms (Fig. 2G; red, bottom-left). Along similar lines, inclusion events in the transition from NP to N (1,391) switched from being the minor (PSI of 21 and 28 in NSC and NP, respectively) to become the major (PSI of 63) isoform in N (Fig. 2G; green, top-right). Vice versa, exclusion events (883) switched from the major (PSI of 66 and 63 in NSC and NP, respectively) to the minor (PSI of 38) isoform in N (Fig. 2G; green, bottom-right). Together, this highlighted progressively consistent patterns of exon inclusion/exclusion in consecutive steps of the neuronal lineage. In addition, and extending on previous observations of exon-inclusion during brain development (Mazin *et al*., 2021; Weyn-Vanhentenryck *et al*., 2018), we also identified a class of N-specific exons virtually absent in NSC that gained inclusion in NP and comprising a considerable proportion (112 out of 281 cassette exons) of microexons (Fig. 2H, top), a class of highly conserved events in the nervous system that almost exclusively show inclusion patterns (Irimia *et al*., 2014; Y. I. Li *et al*., 2015; Mazin *et al*., 2021; Ustianenko *et al*., 2017).

To validate these intriguing observations on inclusion patterns, we next selected 11 events showing: i) low PSI in NSC (< 30), ii) strong PSI gain (≥ 25) in NP and iii) high inclusion level in N (> 75 PSI) and assessed their abundance by qRT-PCR revealing, in all cases, a remarkably high correlation between the bioinformatically predicted PSI and their experimental assessment (R^2^ = 0.95, *p* < 0.0001) (Fig. 2I). Together, these observations reinforce the hypothesis that cell fate specification towards a neurogenic fate involves transcriptome remodeling through AS and, primarily, novel exons inclusion. Most importantly, not only our study highlighted a strong tendency for inclusion events to become the major, if not unique, form in N but also an asymmetry for NP-specific AS patterns to represent a transitory phase prior to the adoption of a definitive, N-specific, AS profile.

### Systematic investigation of protein structural changes resulting from AS

In order to investigate how AS impacts protein structure and, potentially, function we predicted the conformational changes of those isoforms switching during neurogenic commitment using AlphaFold2 (Jumper *et al*., 2021). Modeling of 3D structures and prediction of their global and local conformational changes (Fig. 3A; left) was limited to protein isoforms of up to 800 amino acids, hence, compromising between the computational time needed to model the structures and the quality of the retrieved models. This resulted in the selection of 212 genes of which more than half (127) were associated with GO terms related to organ development, macromolecular complex assembly and protein localization/transport and one third (71) with brain development specifically (not shown). Next, we extracted the 3D structures of the corresponding 987 isoforms undergoing AS during the transitions from NSC to NP and N.

**Figure 3.**
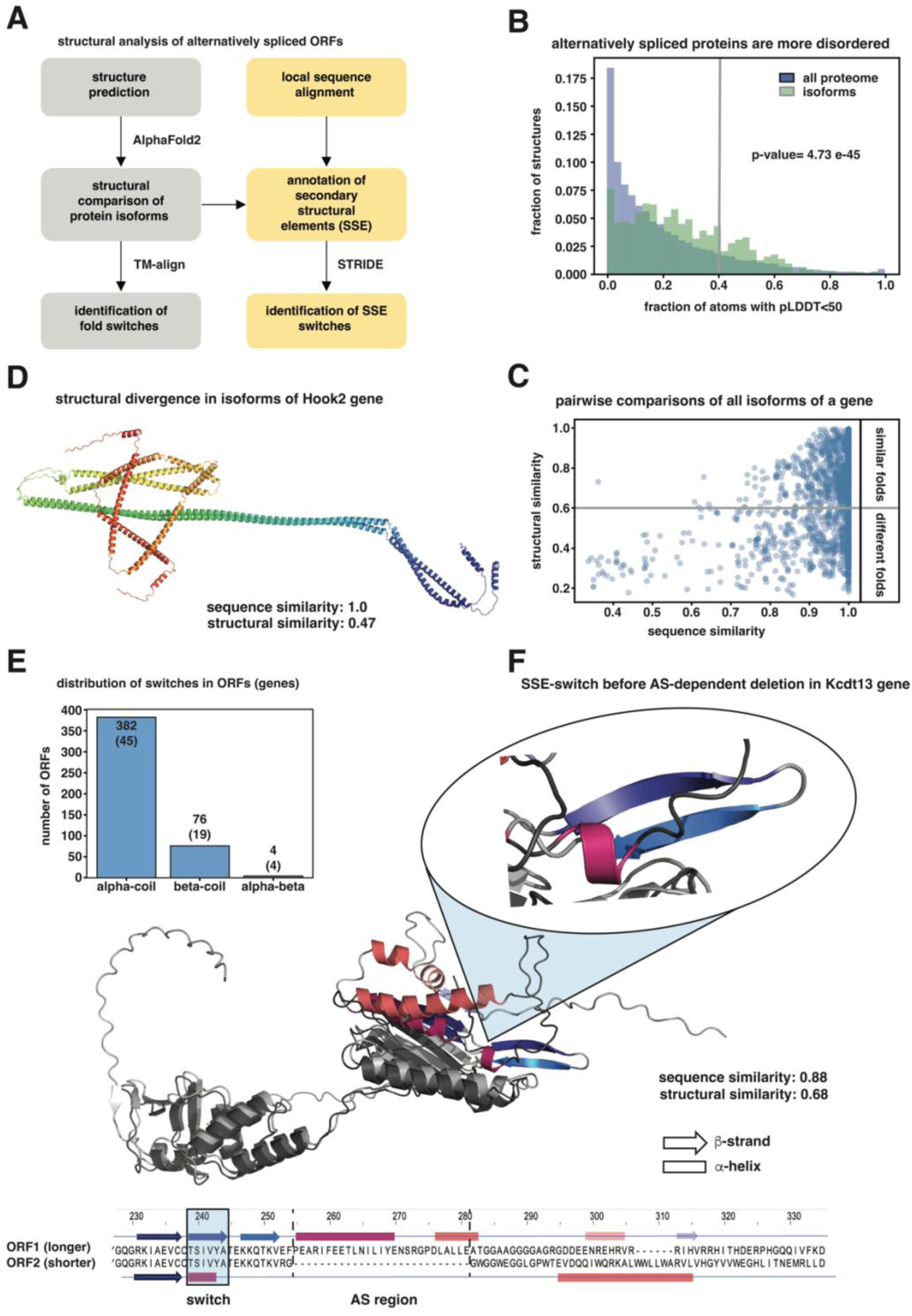
Alternative splicing events induce both global and local conformational changes in 3D structures. **A)** Structural analysis of alternatively spliced proteins including global (grey) or local secondary structure element (SSE) switches (yellow). **B)** Distribution of the fraction of atoms with pLDDT scores < 50 representing disordered structures (threshold indicated by grey bar < 0.4; total structures = 21,615) in our isoform dataset (green) vs. the mouse proteome (blue) (p-value = 4.73e-45, Kolmogorov-Smirnov test). **C)** Pairwise comparisons between all isoforms within one associated gene (TM-score < 0.6 threshold shown as grey line). **D)** Example of global conformational re-arrangement in two isoforms of *Hook2* with high sequence similarity. The change was induced by a small deletion of only two amino acids in a coiled-coli domain. The scores for structural and sequence similarity are shown (scores are normalized by the shorter sequence). **E)** Distribution of the secondary structure element switches in ORFs and associated genes (parentheses). Alpha-coil changes are the most common, but beta-coil and alpha-beta changes were also observed. **F)** Example of an alpha-beta switch in the gene *Kctd13*, encoding a component of a complex required for synaptic transmission. A fragment of the sequence alignment is shown for the two isoforms and corresponding SSEs depicted as rectangles (alpha-helix) or arrows (beta-strand). The C-terminal part of the structures are superimposed (grey and dark grey regions), with the SSE switch highlighted (inset) displaying the two beta-strands (blue and violet) and an alpha-helix (purple) despite identical protein sequences.

In agreement with previous studies (Hegyi *et al*., 2011; Romero *et al*., 2006), we observed that the structures in our set of isoforms were significantly more disordered than the average mouse protein (21,615 proteins from canonical proteome) when using the generally accepted percent threshold of residues with predicted local distance difference test (pLDDT score < 50) (Fig. 3B; gray line). Next, and considering the difficulties in the structural comparison of isoforms with large disordered regions, we limited our analysis to structures where a fraction of residues with very low prediction confidence score (pLDDT score < 50) doesn’t exceed 40%, and compared the sequence and structural similarity of all isoforms associated with one gene.

Not surprisingly, we observed that the template modeling (TM) scores indicative of structural similarity generally increased with increasing sequence similarity (Fig. 3C). However, a remarkably high proportion of genes (78 out of 212; or 37%) mainly with a high sequence similarity (with median equal to 98%) were found to result in isoforms with substantially different structural conformations (TM-score < 0.6) (Fig. 3C; bottom right). As one remarkable example among this group of genes, two *Hook2* isoforms only differed by as little as two amino acids (nearly 100% sequence similarity), yet revealed a remarkably different orientation of their N-terminal helical packing (47% structure similarity) (Fig. 3D). The fact that within this helical packing resides the protein domain essential for *Hook2* function to bind microtubules, implies that such a negligible AS switch by two ammino acids may result in a completely altered function.

While TM-scores are informative of large conformational changes and global structural similarities, we next investigated if AS had the potential to also introduce more subtle, local changes in specific secondary structural elements (SSE). To assess this, we combined SSE annotation with local sequence alignment (Fig. 3A; right) finding primarily alpha- and beta-coil switches and, more rarely, alpha-beta switches (Fig. 3E). Such short sequences that can adopt both alpha and beta conformations depending on the structural and sequence context within which they are embedded are known as chameleon sequences (W. Li *et al*., 2015). In our set of genes, we found that these chameleon transitions could be induced just by the presence or absence of a distant spliced region with the rest of the structural and sequence neighborhood context of the protein being nearly identical. A notable example of an alpha-beta switch was found in the gene *Kctd13* (Fig. 3F), encoding a component of a complex required for synaptic transmission (Escamilla *et al*., 2017) and implicated in neurodevelopmental disorders such as macrocephaly (Golzio *et al*., 2012) (although a subsequent study could not confirm the association with this latter phenotype (Escamilla *et al*., 2017)). This example highlights how identical sequences can adopt different structural conformations as a result of an AS event occurring 10 amino acids away.

Several other examples of both global and local conformational changes resulting from AS events occurring within or outside the affected protein domain were found in our study (Supplementary Material). It will be intriguing in future studies to investigate how many of them can lead to protein functional diversification relevant for cell fate commitment during brain development.

## DISCUSSION

Our study explores a new avenue for assessing gene function in cell fate commitment by looking at the potential diversification of protein structure resulting from AS rather than, as universally adopted, measuring gene expression alone. As a traditionally poorly investigated aspect of cell biology, a comprehensive assessment of AS required the advent of new sequencing technologies and bioinformatic tools. Similarly, reliable prediction of protein structures from their sequence alone required the implementation of machine learning to interrogate depository databases of crystal structures. Here, we combined these tools starting with a validated reporter mouse model allowing the identification of cell populations recapitulating neurogenic commitment during brain development.

In doing so, our work advances the field in several contexts. First, it describes a novel methodological approach to assess AS by the combination of SRS and LRS and further extend this to predict its effects on protein structure by machine learning. Second, it provides a resource for the field as new transcriptome annotation of the developing mouse brain including nearly 50,000 novel isoforms and 2,500 previously unknown splicing events. Third, it extends previous reports on the significance of AS in the context of neuronal maturation (Liu *et al*., 2018; Zhang *et al*., 2016) to also reveal its full extent and potential significance in cell fate commitment of stem and progenitor cells. Several aspects of our study are worth discussing toward better understanding the biological significance of our observations.

In agreement with previous assessments of AS profiles during organogenesis (Mazin *et al*., 2021), our analysis in the context of cortical development revealed patterns of AS events strongly skewed towards inclusion of exons in both transitions from NSC to NP and NP to N. Most importantly, the magnitude of the observed inclusion events was such that, on average, isoforms only marginally represented in NSC, if any, became the main, if not unique, isoform in N with NP representing a transient intermediate cellular state between the two. In essence, AS alone was revealed to have a much greater impact in remodeling the transcriptome profile from NSC to N than previously thought and independently from changes in gene expression.

Also expanding on previous findings on a peculiar class of short exons in the nervous system (Irimia *et al*., 2014; Y. I. Li *et al*., 2015; Mazin *et al*., 2021; Ustianenko *et al*., 2017), we found that virtually all alternatively spliced microexons were progressively more included as development proceeded from NSC to N and accounting for more than one third (40%) of included cassette exons during the neurogenic commitment step. This observation strengthens the hypothesis that this class of highly-conserved exons might have a central role in redefining neuronal identity highlighting the importance in our study of assessing specific cell sub-populations to reveal the full potential of AS patterns in fate-commitment.

Next, we investigated the impact of AS on protein structure by AlphaFold2 finding a high proportion of disordered regions in our set of alternatively spliced genes. In fact, AS often affects intrinsically disordered regions (W. Li et al., 2015) and tends to avoid globular domains or affect them only marginally at locations where the exposed hydrophobic surface is minimal (Hegyi *et al*., 2011). An emerging concept in the field is that differential inclusion of disordered segments can favor new protein interactions and, hence, change the context in which the molecular function of the protein is carried out (Buljan *et al*., 2013). Our study supports this notion with a significant increase in disordered isoforms arising concomitantly with neurogenic commitment.

Finally, it is important to bear in mind that while AlphaFold2 has remarkable accuracy in predicting ordered proteins conserved across evolution, its accuracy significantly decreases when predicting disordered regions and sequences with only few homologs. Clearly, applying stringent quality filters and limiting our analysis to the comparison of 3D conformations with less or equal to 40% disordered regions would reduce, but certainly not remove, the occurrence of noise in predicted conformations due to uncertainty that are not biologically relevant. Still, while false results are difficult to quantify, the extent and number of the structural changes observed between isoforms that were nearly identical in sequence appeared to be far greater than what prediction errors seems to justify. This was similarly the case both when the structural change occurred within the AS event as well, more remarkably, when the event was far away. In addition, we also found several chameleon sequences that adopted different secondary structures in specific cell types as a result of AS. While these regions are long known to exist, their structural switch was assumed to be dependent on substantial changes in their structural and sequence contexts (Gendoo and Harrison, 2011; W. Li *et al*., 2015) as opposed to, as observed in our study, being triggered by small perturbations within nearly identical sequence contexts.

While more studies are need to validate the predicted structural changes, their impact on protein function and, most importantly, their relevance for brain development, we hope that our study may provide a new resource and conceptual framework to better address the significance of AS during cell fate commitment and organogenesis.

## MATERIALS AND METHODS

### SRS and LRS transcriptome assembly of NSC, NP and N

A previous SRS dataset of FAC-sorted *Btg2*::RFP/*Tubb3*::GFP cells of the E14.5 mouse cortex obtained in three biological replicates (Aprea *et al*., 2015) was used to generate a new transcriptome assembly identifying NSC, NP and N. Paired-end reads were mapped to the Ensembl (GRCm38.p6) mouse genome using Hisat2 (Kim *et al*., 2019) and Stringtie used to assemble the mapped reads into transcripts using Ensembl as a reference (only transcripts with support level 1/2) (Pertea *et al*., 2015). Only transcripts with > 10 supporting reads were included and fusion transcripts removed. Reference transcriptome was not included in the final assembly. For the generation of a new LRS dataset, Isoseq library preparation was performed on 300 ng of RNA (RIN ≥ 9.3) of each cell type (isolated as described above) using NEBNext® Single Cell/Low Input cDNA Synthesis & Amplification (NEB), Iso-Seq Express Oligo Kit (PacBio) and SMRTbell Express Template Prep Kit 2.0 (PacBio). Samples were sequenced using PacBio Sequel II 8M SMRT cell (version 2.1) and data processed with Isoseq3 workflow (≥ 3 passes and accuracy ≥ 0.9 for CCS reads and ≥ 7 passes for high-quality transcripts). High-quality transcripts were aligned to M. musculus GRCm38/mm10, collapsed to non-redundant transcripts, and 5’ degraded transcripts and potential artifacts removed using either pipeline 1 (including gmap (Wu and Watanabe, 2005), cDNA_Cupcake (https://github.com/Magdoll/cDNA_Cupcake) and Sqanti3 (Tardaguila *et al*., 2018)) or pipeline 2 (including deSALT (Liu *et al*., 2019), Tama (Kuo *et al*., 2020) and Sqanti3)). Transcripts from pipelines 1 and 2, excluding fusion transcripts, were merged into a LRS transcriptome assembly with Tama merge using a tolerance of 25 nt for the 5’ ends (-a 25 parameter) and merged with the SRS transcriptome in a unique assembly. Sqanti3 was used for classification of transcripts by comparing it with the best-matching reference transcript from Ensembl (GRCm38.p6), NCBI_RefSeq, and Gencode (vM10). Transcripts were considered known when containing only known splice as either all (full-splice match) or only part (incomplete splice-match) of the best-matching reference. Transcripts were classified as novel in catalog (or not in catalog) if containing a novel combination of known donor and acceptor sites (or novel donor and/or acceptor site).

All experimental procedures were performed according to local regulations and approved by the “Landesdirektion Sachsen” under the licenses 11-1-2011-41 and TVV 16/2018.

### Detection of splicing events

Paired-end reads (Aprea *et al*., 2015) were mapped to the mouse genome annotated with the developing cortex transcriptome assembly using STAR (Dobin *et al*., 2013). Whippet (Sterne-Weiler *et al*., 2018) was then used to estimate exon and intron inclusion in each replicate. Lowly expressed genes (< 0.5 transcripts per million) were excluded from analysis. AS events were defined as absolute PSI change ≥ 10 with a minimum probability of 0.9 and a confidence interval width ≤ 0.3. When the exon boundaries did not coincide among a genés transcripts (alternative 5’ or 3’ splice site choice), Whippet divide the exon into nodes and calculate a PSI for each of them. Contiguous nodes showing similar PSI levels and patterns and likely belonging to a single exon were fused, their PSI averaged and considered a single event. The cluster module of DAVID (Huang *et al*., 2009; Sherman *et al*., 2022) was used for clustering and enrichment evaluation of gene ontology terms of differentially spliced genes. Multiexonic genes expressed in the cell populations of interest were used as a background.

### Identification and validation of AS isoforms and novel splicing events

Isoforms with high abundance (> 4 unique splice-junction reads across all splice-junctions) were considered and AS events coordinates compared to the transcripts annotated in our dataset. Events were assigned to an inclusion isoform if their coordinates overlapped, at least partially, with an exon or to an exclusion isoform if they were located within an intron. AS events without a corresponding inclusion or exclusion isoform were assigned to an Ensembl or NCBI_RefSeq isoform using the criteria above. Only AS events assigned to at least one inclusion and one exclusion isoform were considered for further analysis. AS events and exclusion sites predicted to have a novel acceptor and/or donor site or a new combination of known splice sites were identified by Sqanti3, while the AS events containing novel sequences were identified by bedtools. For novel isoforms and AS events validation, RNA with RIN > 8.0 from C57BL/6J lateral cortices was collected and qRT-PCR performed to assess novel splice junctions, as well as levels of inclusion changes of AS events. For each target sequence, amplification by PCR was carried out with 1 ng of cDNA with standard conditions: 98 °C 30 sec, 27-33 cycles of 98°C 15 sec, 68°C 25 sec, 72°C 30 sec, 72°C 2 min. PCR products were resolved in 15% polyacrylamide gels.

### Structural analysis

AlphaFold2 was used to model the 3D structures of isoforms (Jumper *et al*., 2021). To access global structural rearrangements between isoforms, global structural alignment was performed between each pair of isoforms within each gene using the TM-align algorithm (Zhang and Skolnick, 2005) computing the TM-score for each alignment and allowing the assessment of structural similarity between proteins. To identify local structural rearrangements, a pipeline was developed (Fig. 3A) including protein local sequence alignment by the Smith-Waterman algorithm (python package Biopython). The computed AlphaFold structural models were annotated with secondary structure elements (SSE) by STRIDE (Heinig and Frishman, 2004). SSE switching regions of isoforms were defined when changes in the assignment of SSE for the corresponding regions in protein sequence (identified by alignment) were observed for ≥ 5 consecutive amino acids. Figures were generated by Pymol, Jalview and Python matplotlib.

## REFERENCES

1. An, D., Cao, H., Li, C., Humbeck, K. and Wang, W. 2018. Isoform Sequencing and State-of-Art Applications for Unravelling Complexity of Plant Transcriptomes. Genes, 9: 43.

2. Aprea, J., Lesche, M., Massalini, S., Prenninger, S., Alexopoulou, D., Dahl, A., Hiller, M. and Calegari, F. 2015. Identification and expression patterns of novel long non-coding RNAs in neural progenitors of the developing mammalian cortex. Neurogenesis, 2: e995524.

3. Aprea, J., Prenninger, S., Dori, M., Ghosh, T., Monasor, L.S., Wessendorf, E., Zocher, S., Massalini, S., Alexopoulou, D., Lesche, M., Dahl, A., Groszer, M., Hiller, M. and Calegari, F. 2013. Transcriptome sequencing during mouse brain development identifies long non-coding RNAs functionally involved in neurogenic commitment: LncRNAs control neurogenesis. The EMBO Journal, 32: 3145–3160.

4. Artegiani, B., De Jesus Domingues, A.M., Bragado Alonso, S., Brandl, E., Massalini, S., Dahl, A. and Calegari, F. 2015. Tox: a multifunctional transcription factor and novel regulator of mammalian corticogenesis. The EMBO Journal, 34: 896–910.

5. Arzalluz-Luque, Á. and Conesa, A. 2018. Single-cell RNAseq for the study of isoforms—how is that possible? Genome Biol, 19: 110.

6. Barbosa-Morais, N.L., Irimia, M., Pan, Q., Xiong, H.Y., Gueroussov, S., Lee, L.J., Slobodeniuc, V., Kutter, C., Watt, S., Çolak, R., Kim, T., Misquitta-Ali, C.M., Wilson, M.D., Kim, P.M., Odom, D.T., Frey, B.J. and Blencowe, B.J. 2012. The Evolutionary Landscape of Alternative Splicing in Vertebrate Species. 338: 8.

7. Buen Abad Najar, C.F., Yosef, N. and Lareau, L.F. 2020. Coverage-dependent bias creates the appearance of binary splicing in single cells. eLife, 9: e54603.

8. Buljan, M., Chalancon, G., Dunker, A.K., Bateman, A., Balaji, S., Fuxreiter, M. and Babu, M.M. 2013. Alternative splicing of intrinsically disordered regions and rewiring of protein interactions. Current Opinion in Structural Biology, 23: 443–450.

9. Buljan, M., Chalancon, G., Eustermann, S., Wagner, G.P., Fuxreiter, M., Bateman, A. and Babu, M.M. 2012. Tissue-Specific Splicing of Disordered Segments that Embed Binding Motifs Rewires Protein Interaction Networks. Molecular Cell, 46: 871–883.

10. Chau, K.K., Zhang, P., Urresti, J., Amar, M., Pramod, A.B., Chen, J., Thomas, A., Corominas, R., Lin, G.N. and Iakoucheva, L.M. 2021. Full-length isoform transcriptome of the developing human brain provides further insights into autism. Cell Reports, 36: 109631.

11. Chen, L., Bush, S.J., Tovar-Corona, J.M., Castillo-Morales, A. and Urrutia, A.O. 2014. Correcting for Differential Transcript Coverage Reveals a Strong Relationship between Alternative Splicing and Organism Complexity. Molecular Biology and Evolution, 31: 1402–1413.

12. Cunningham, F., Allen, J.E., Allen, J., Alvarez-Jarreta, J., Amode, M.R., Armean, I.M., Austine-Orimoloye, O., Azov, A.G., Barnes, I., Bennett, R., Berry, A., Bhai, J., Bignell, A., Billis, K., Boddu, S., Brooks, L., Charkhchi, M., Cummins, C., Da Rin Fioretto, L., et al. 2022. Ensembl 2022. Nucleic Acids Research, 50: D988–D995.

13. Dobin, A., Davis, C.A., Schlesinger, F., Drenkow, J., Zaleski, C., Jha, S., Batut, P., Chaisson, M. and Gingeras, T.R. 2013. STAR: ultrafast universal RNA-seq aligner. Bioinformatics, 29: 15–21.

14. Dori, M., Alieh, L.H.A., Cavalli, D., Massalini, S., Lesche, M., Dahl, A. and Calegari, F. 2019. Sequence and expression levels of circular RNAs in progenitor cell types during mouse corticogenesis. Life Sci. Alliance, 2: 10.

15. Dori, M., Cavalli, D., Lesche, M., Massalini, S., Alieh, L.H.A., de Toledo, B.C., Khudayberdiev, S., Schratt, G., Dahl, A. and Calegari, F. 2020. MicroRNA profiling of mouse cortical progenitors and neurons reveals miR-486-5p as a regulator of neurogenesis. Development, 147: 8.

16. Ellis, J.D., Barrios-Rodiles, M., Çolak, R., Irimia, M., Kim, T., Calarco, J.A., Wang, X., Pan, Q., O’Hanlon, D., Kim, P.M., Wrana, J.L. and Blencowe, B.J. 2012. Tissue-Specific Alternative Splicing Remodels Protein-Protein Interaction Networks. Molecular Cell, 46: 884–892.

17. Escamilla, C.O., Filonova, I., Walker, A.K., Xuan, Z.X., Holehonnur, R., Espinosa, F., Liu, S., Thyme, S.B., López-García, I.A., Mendoza, D.B., Usui, N., Ellegood, J., Eisch, A.J., Konopka, G., Lerch, J.P., Schier, A.F., Speed, H.E. and Powell, C.M. 2017. Kctd13 deletion reduces synaptic transmission via increased RhoA. Nature, 551: 227–231.

18. Florio, M., Namba, T., Pääbo, S., Hiller, M. and Huttner, W.B. 2016. A single splice site mutation in human-specific *ARHGAP11B* causes basal progenitor amplification. Sci. Adv., 2: e1601941.

19. Furlanis, E. and Scheiffele, P. 2018. Regulation of Neuronal Differentiation, Function, and Plasticity by Alternative Splicing. Annu. Rev. Cell Dev. Biol., 34: 451–469.

20. Gendoo, D.M.A. and Harrison, P.M. 2011. Discordant and chameleon sequences: Their distribution and implications for amyloidogenicity. Protein Science, 20: 567–579.

21. Golzio, C., Willer, J., Talkowski, M.E., Oh, E.C., Taniguchi, Y., Jacquemont, S., Reymond, A., Sun, M., Sawa, A., Gusella, J.F., Kamiya, A., Beckmann, J.S. and Katsanis, N. 2012. KCTD13 is a major driver of mirrored neuroanatomical phenotypes of the 16p11.2 copy number variant. Nature, 485: 363–367.

22. Gupta, I., Collier, P.G., Haase, B., Mahfouz, A., Joglekar, A., Floyd, T., Koopmans, F., Barres, B., Smit, A.B., Sloan, S.A., Luo, W., Fedrigo, O., Ross, M.E. and Tilgner, H.U. 2018. Single-cell isoform RNA sequencing characterizes isoforms in thousands of cerebellar cells. Nat Biotechnol, 36: 1197–1202.

23. Hahn, M.W. and Wray, G.A. 2002. The g-value paradox. Evol Dev, 4: 73–75.

24. Han, H., Best, A.J., Braunschweig, U., Mikolajewicz, N., Li, J.D., Roth, J., Chowdhury, F., Mantica, F., Nabeel-Shah, S., Parada, G., Brown, K.R., O’Hanlon, D., Wei, J., Yao, Y., Zid, A.A., Comsa, L.C., Jen, M., Wang, J., Datti, A., et al. 2022. Systematic exploration of dynamic splicing networks reveals conserved multistage regulators of neurogenesis. Molecular Cell, 82: 2982–2999.e14.

25. Hegyi, H., Kalmar, L., Horvath, T. and Tompa, P. 2011. Verification of alternative splicing variants based on domain integrity, truncation length and intrinsic protein disorder. Nucleic Acids Research, 39: 1208–1219.

26. Heinig, M. and Frishman, D. 2004. STRIDE: a web server for secondary structure assignment from known atomic coordinates of proteins. Nucleic Acids Research, 32: W500–W502.

27. Huang, D.W., Sherman, B.T. and Lempicki, R.A. 2009. Systematic and integrative analysis of large gene lists using DAVID bioinformatics resources. Nat Protoc, 4: 44–57.

28. Irimia, M., Weatheritt, R.J., Ellis, J.D., Parikshak, N.N., Gonatopoulos-Pournatzis, T., Babor, M., Quesnel-Vallières, M., Tapial, J., Raj, B., O’Hanlon, D., Barrios-Rodiles, M., Sternberg, M.J.E., Cordes, S.P., Roth, F.P., Wrana, J.L., Geschwind, D.H. and Blencowe, B.J. 2014. A Highly Conserved Program of Neuronal Microexons Is Misregulated in Autistic Brains. Cell, 159: 1511– 1523.

29. Joglekar, A., Prjibelski, A., Mahfouz, A., Collier, P., Lin, S., Schlusche, A.K., Marrocco, J., Williams, S.R., Haase, B., Hayes, A., Chew, J.G., Weisenfeld, N.I., Wong, M.Y., Stein, A.N., Hardwick, S.A., Hunt, T., Wang, Q., Dieterich, C., Bent, Z., et al. 2021. A spatially resolved brain region- and cell type-specific isoform atlas of the postnatal mouse brain. Nat Commun, 12: 463.

30. Jumper, J., Evans, R., Pritzel, A., Green, T., Figurnov, M., Ronneberger, O., Tunyasuvunakool, K., Bates, R., Žídek, A., Potapenko, A., Bridgland, A., Meyer, C., Kohl, S.A.A., Ballard, A.J., Cowie, A., Romera-Paredes, B., Nikolov, S., Jain, R., Adler, J., et al. 2021. Highly accurate protein structure prediction with AlphaFold. Nature, 596: 583–589.

31. Kelemen, O., Convertini, P., Zhang, Z., Wen, Y., Shen, M., Falaleeva, M. and Stamm, S. 2013. Function of alternative splicing. Gene, 514: 1–30.

32. Kim, D., Paggi, J.M., Park, C., Bennett, C. and Salzberg, S.L. 2019. Graph-based genome alignment and genotyping with HISAT2 and HISAT-genotype. Nat Biotechnol, 37: 907–915.

33. Kuo, R.I., Cheng, Y., Zhang, R., Brown, J.W.S., Smith, J., Archibald, A.L. and Burt, D.W. 2020. Illuminating the dark side of the human transcriptome with long read transcript sequencing. BMC Genomics, 21: 751.

34. Leung, S.K., Jeffries, A.R., Castanho, I., Jordan, B.T., Moore, K., Davies, J.P., Dempster, E.L., Bray, N.J., O’Neill, P., Tseng, E., Ahmed, Z., Collier, D.A., Jeffery, E.D., Prabhakar, S., Schalkwyk, L., Jops, C., Gandal, M.J., Sheynkman, G.M., Hannon, E., et al. 2021. Full-length transcript sequencing of human and mouse cerebral cortex identifies widespread isoform diversity and alternative splicing. Cell Reports, 37: 110022.

35. Li, W., Kinch, L.N., Karplus, P.A. and Grishin, N.V. 2015. CHSEQ : A database of chameleon sequences. Protein Science, 24: 1075–1086.

36. Li, Y.I., Sanchez-Pulido, L., Haerty, W. and Ponting, C.P. 2015. RBFOX and PTBP1 proteins regulate the alternative splicing of micro-exons in human brain transcripts. Genome Res., 25: 1–13.

37. Liu, B., Liu, Y., Li, J., Guo, H., Zang, T. and Wang, Y. 2019. deSALT: fast and accurate long transcriptomic read alignment with de Bruijn graph-based index. Genome Biol, 20: 274.

38. Liu, J., Geng, A., Wu, X., Lin, R.-J. and Lu, Q. 2018. Alternative RNA Splicing Associated With Mammalian Neuronal Differentiation. Cerebral Cortex, 28: 2810–2816.

39. Marasco, L.E. and Kornblihtt, A.R. 2023. The physiology of alternative splicing. Nat Rev Mol Cell Biol, 24: 242–254.

40. Mazin, P.V., Khaitovich, P., Cardoso-Moreira, M. and Kaessmann, H. 2021. Alternative splicing during mammalian organ development. Nat Genet, 53: 925–934.

41. Mestres, I. and Calegari, F. 2023. 4931414P19Rik, a microglia chemoattractant secreted by neural progenitors, modulates neuronal migration during corticogenesis. Development, 150: dev201574.

42. Morillon, A. and Gautheret, D. 2019. Bridging the gap between reference and real transcriptomes. Genome Biol, 20: 112.

43. Nilsen, T.W. and Graveley, B.R. 2010. Expansion of the eukaryotic proteome by alternative splicing. Nature, 463: 457–463.

44. Noack, F., Pataskar, A., Schneider, M., Buchholz, F., Tiwari, V.K. and Calegari, F. 2019. Assessment and site-specific manipulation of DNA (hydroxy-)methylation during mouse corticogenesis. Life Sci. Alliance, 2: e201900331.

45. Oikonomopoulos, S., Bayega, A., Fahiminiya, S., Djambazian, H., Berube, P. and Ragoussis, J. 2020. Methodologies for Transcript Profiling Using Long-Read Technologies. Front. Genet., 11: 606.

46. Pertea, M., Pertea, G.M., Antonescu, C.M., Chang, T.-C., Mendell, J.T. and Salzberg, S.L. 2015. StringTie enables improved reconstruction of a transcriptome from RNA-seq reads. Nat Biotechnol, 33: 290–295.

47. Pickrell, J.K., Pai, A.A., Gilad, Y. and Pritchard, J.K. 2010. Noisy Splicing Drives mRNA Isoform Diversity in Human Cells. PLoS Genet, 6: e1001236.

48. Quesnel-Vallières, M., Dargaei, Z., Irimia, M., Gonatopoulos-Pournatzis, T., Ip, J.Y., Wu, M., Sterne-Weiler, T., Nakagawa, S., Woodin, M.A., Blencowe, B.J. and Cordes, S.P. 2016. Misregulation of an Activity-Dependent Splicing Network as a Common Mechanism Underlying Autism Spectrum Disorders. Molecular Cell, 64: 1023–1034.

49. Raj, B. and Blencowe, B.J. 2015. Alternative Splicing in the Mammalian Nervous System: Recent Insights into Mechanisms and Functional Roles. Neuron, 87: 14–27.

50. Reixachs-Solé, M. and Eyras, E. 2022. Uncovering the impacts of alternative splicing on the proteome with current omics techniques. WIREs RNA, 13.

51. Rodriguez, J.M. and Pozo, F. 2020. An analysis of tissue-specific alternative splicing at the protein level. PLoS Comput Biol, 16: e1008287.

52. Romero, P.R., Zaidi, S., Fang, Y.Y., Uversky, V.N., Radivojac, P., Oldfield, C.J., Cortese, M.S., Sickmeier, M., LeGall, T., Obradovic, Z. and Dunker, A.K. 2006. Alternative splicing in concert with protein intrinsic disorder enables increased functional diversity in multicellular organisms. Proc Natl Acad Sci USA, 103: 6.

53. Sahu, S.K., Agirre, E., Inayatullah, M., Mahesh, A., Tiwari, N., Lavin, D.P., Singh, A., Strand, S., Diken, M., Luco, R.F., Belmonte, J.C.I. and Tiwari, V.K. 2022. A complex epigenome-splicing crosstalk governs epithelial-to-mesenchymal transition in metastasis and brain development. Nat Cell Biol, 24: 1265–1277.

54. Sarantopoulou, D., Brooks, T.G., Nayak, S., Mrčela, A., Lahens, N.F. and Grant, G.R. 2021. Comparative evaluation of full-length isoform quantification from RNA-Seq. BMC Bioinformatics, 22: 266.

55. Saudemont, B., Popa, A., Parmley, J.L., Rocher, V., Blugeon, C., Necsulea, A., Meyer, E. and Duret, L. 2017. The fitness cost of mis-splicing is the main determinant of alternative splicing patterns. Genome Biol, 18: 208.

56. Schaefke, B., Sun, W., Li, Y., Fang, L. and Chen, W. 2018. The evolution of posttranscriptional regulation. WIREs RNA, 9: e1485.

57. Sherman, B.T., Hao, M., Qiu, J., Jiao, X., Baseler, M.W., Lane, H.C., Imamichi, T. and Chang, W. 2022. DAVID: a web server for functional enrichment analysis and functional annotation of gene lists (2021 update). Nucleic Acids Research, 50: W216–W221.

58. Shumate, A., Wong, B., Pertea, G. and Pertea, M. 2022. Improved transcriptome assembly using a hybrid of long and short reads with StringTie. PLoS Comput Biol, 18: e1009730.

59. Sterne-Weiler, T., Weatheritt, R.J., Best, A.J., Ha, K.C.H. and Blencowe, B.J. 2018. Efficient and Accurate Quantitative Profiling of Alternative Splicing Patterns of Any Complexity on a Laptop. Molecular Cell, 72: 187–200.e6.

60. Tardaguila, M., de la Fuente, L., Marti, C., Pereira, C., Pardo-Palacios, F.J., del Risco, H., Ferrell, M., Mellado, M., Macchietto, M., Verheggen, K., Edelmann, M., Ezkurdia, I., Vazquez, J., Tress, M., Mortazavi, A., Martens, L., Rodriguez-Navarro, S., Moreno-Manzano, V. and Conesa, A. 2018. SQANTI: extensive characterization of long-read transcript sequences for quality control in full-length transcriptome identification and quantification. Genome Res., 28: 396–411.

61. Thomas, C.A. 1971. The genetic organization of chromosomes. Annu. Rev. Genet., 5: 237–256.

62. Tress, M.L., Abascal, F. and Valencia, A. 2017. Alternative Splicing May Not Be the Key to Proteome Complexity. Trends in Biochemical Sciences, 42: 98–110.

63. Ustianenko, D., Weyn-Vanhentenryck, S.M. and Zhang, C. 2017. Microexons: discovery, regulation, and function: Microexons: discovery, regulation, and function. WIREs RNA, 8: e1418.

64. Varadi, M., Anyango, S., Deshpande, M., Nair, S., Natassia, C., Yordanova, G., Yuan, D., Stroe, O., Wood, G., Laydon, A., Žídek, A., Green, T., Tunyasuvunakool, K., Petersen, S., Jumper, J., Clancy, E., Green, R., Vora, A., Lutfi, M., et al. 2022. AlphaFold Protein Structure Database: massively expanding the structural coverage of protein-sequence space with high-accuracy models. Nucleic Acids Research, 50: D439–D444.

65. Veiga, D.F.T., Nesta, A., Zhao, Y., Mays, A.D., Huynh, R., Rossi, R., Wu, T.-C., Palucka, K., Anczukow, O., Beck, C.R. and Banchereau, J. 2022. A comprehensive long-read isoform analysis platform and sequencing resource for breast cancer. Sci. Adv., 8: eabg6711.

66. Verdile, V., Riccioni, V., Guerra, M., Ferrante, G., Sette, C., Valle, C., Ferri, A. and Paronetto, M.P. 2023. An impaired splicing program underlies differentiation defects in hSOD1G93A neural progenitor cells. Cell. Mol. Life Sci., 80: 236.

67. Verta, J.-P. and Jacobs, A. 2022. The role of alternative splicing in adaptation and evolution. Trends in Ecology & Evolution, 37: 299–308.

68. Villalba, A., Götz, M. and Borrell, V. 2021. The regulation of cortical neurogenesis. In: Current Topics in Developmental Biology, pp. 1–66.

69. Westoby, J., Artemov, P., Hemberg, M. and Ferguson-Smith, A. 2020. Obstacles to detecting isoforms using full-length scRNA-seq data. Genome Biol, 21: 74.

70. Weyn-Vanhentenryck, S.M., Feng, H., Ustianenko, D., Duffié, R., Yan, Q., Jacko, M., Martinez, J.C., Goodwin, M., Zhang, X., Hengst, U., Lomvardas, S., Swanson, M.S. and Zhang, C. 2018. Precise temporal regulation of alternative splicing during neural development. Nat Commun, 9: 2189.

71. Wu, T.D. and Watanabe, C.K. 2005. GMAP: a genomic mapping and alignment program for mRNA and EST sequences. Bioinformatics, 21: 1859–1875.

72. Yang, P., Wang, D. and Kang, L. 2021. Alternative splicing level related to intron size and organism complexity. BMC Genomics, 22: 853.

73. Yeo, G., Holste, D., Kreiman, G. and Burge, C.B. 2004. Variation in alternative splicing across human tissues. Genome Biol, 5: R74.

74. Zhang, D., Guelfi, S., Garcia-Ruiz, S., Costa, B., Reynolds, R.H., D’Sa, K., Liu, W., Courtin, T., Peterson, A., Jaffe, A.E., Hardy, J., Botía, J.A., Collado-Torres, L. and Ryten, M. 2020. Incomplete annotation has a disproportionate impact on our understanding of Mendelian and complex neurogenetic disorders. Sci. Adv., 6: eaay8299.

75. Zhang, X., Chen, M.H., Wu, X., Kodani, A., Fan, J., Doan, R., Ozawa, M., Ma, J., Yoshida, N., Reiter, J.F., Black, D.L., Kharchenko, P.V., Sharp, P.A. and Walsh, C.A. 2016. Cell-Type-Specific Alternative Splicing Governs Cell Fate in the Developing Cerebral Cortex. Cell, 166: 1147–1162.e15.

76. Zhang, Y. and Skolnick, J. 2005. TM-align: a protein structure alignment algorithm based on the TM-score. Nucleic Acids Research, 33: 2302–2309.

